## supplementary figures for "De novo annotation of the wheat pan-genome reveals complexity and diversity of the hexaploid wheat pan-transcriptome"

### Slide 1
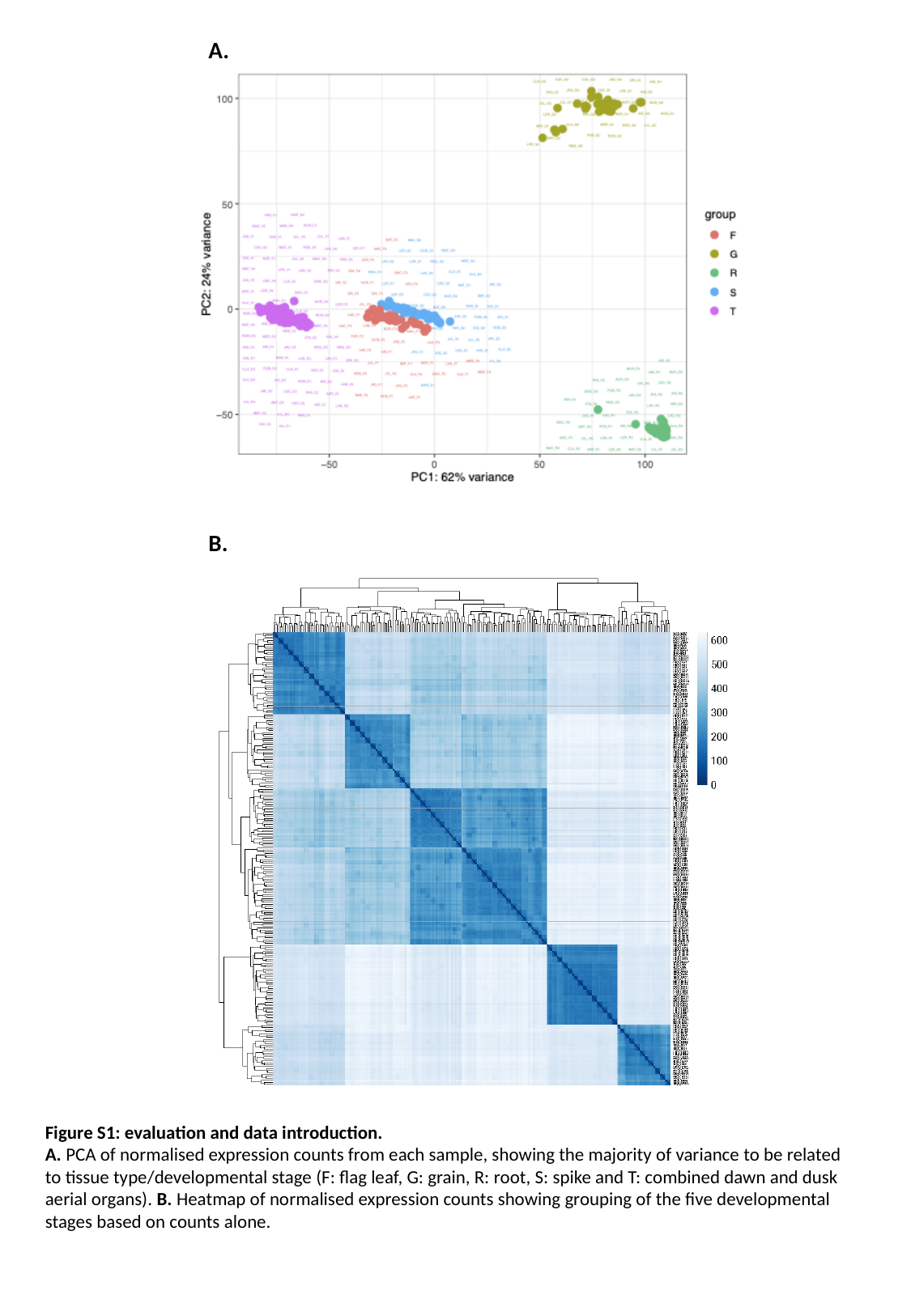

A.
B.
Figure S1: evaluation and data introduction.
A. PCA of normalised expression counts from each sample, showing the majority of variance to be related to tissue type/developmental stage (F: flag leaf, G: grain, R: root, S: spike and T: combined dawn and dusk aerial organs). B. Heatmap of normalised expression counts showing grouping of the five developmental stages based on counts alone.

### Slide 2
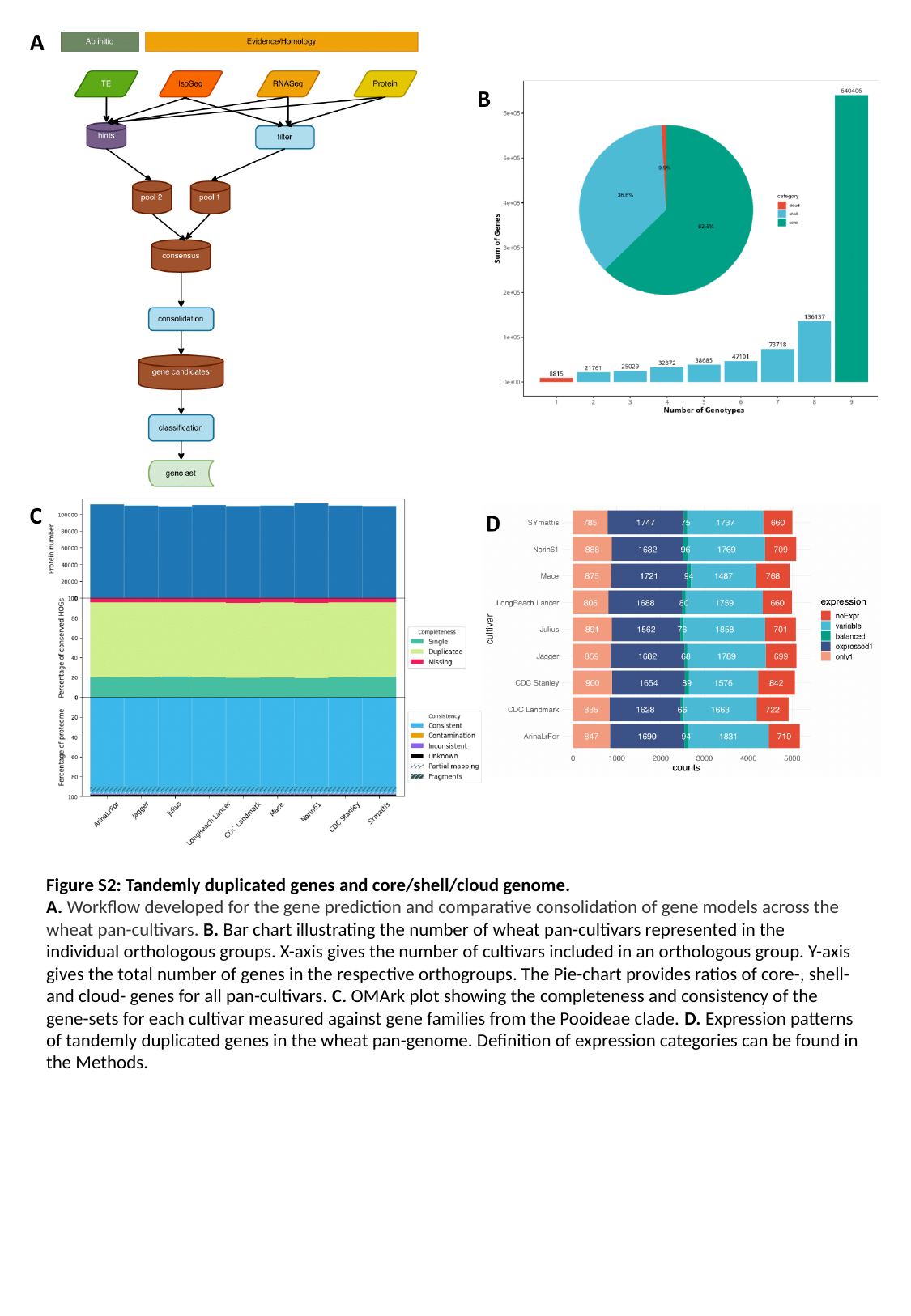

Figure S2: Tandemly duplicated genes and core/shell/cloud genome.
A. Workflow developed for the gene prediction and comparative consolidation of gene models across the wheat pan-cultivars. B. Bar chart illustrating the number of wheat pan-cultivars represented in the individual orthologous groups. X-axis gives the number of cultivars included in an orthologous group. Y-axis gives the total number of genes in the respective orthogroups. The Pie-chart provides ratios of core-, shell- and cloud- genes for all pan-cultivars. C. OMArk plot showing the completeness and consistency of the gene-sets for each cultivar measured against gene families from the Pooideae clade. D. Expression patterns of tandemly duplicated genes in the wheat pan-genome. Definition of expression categories can be found in the Methods.

### Slide 3
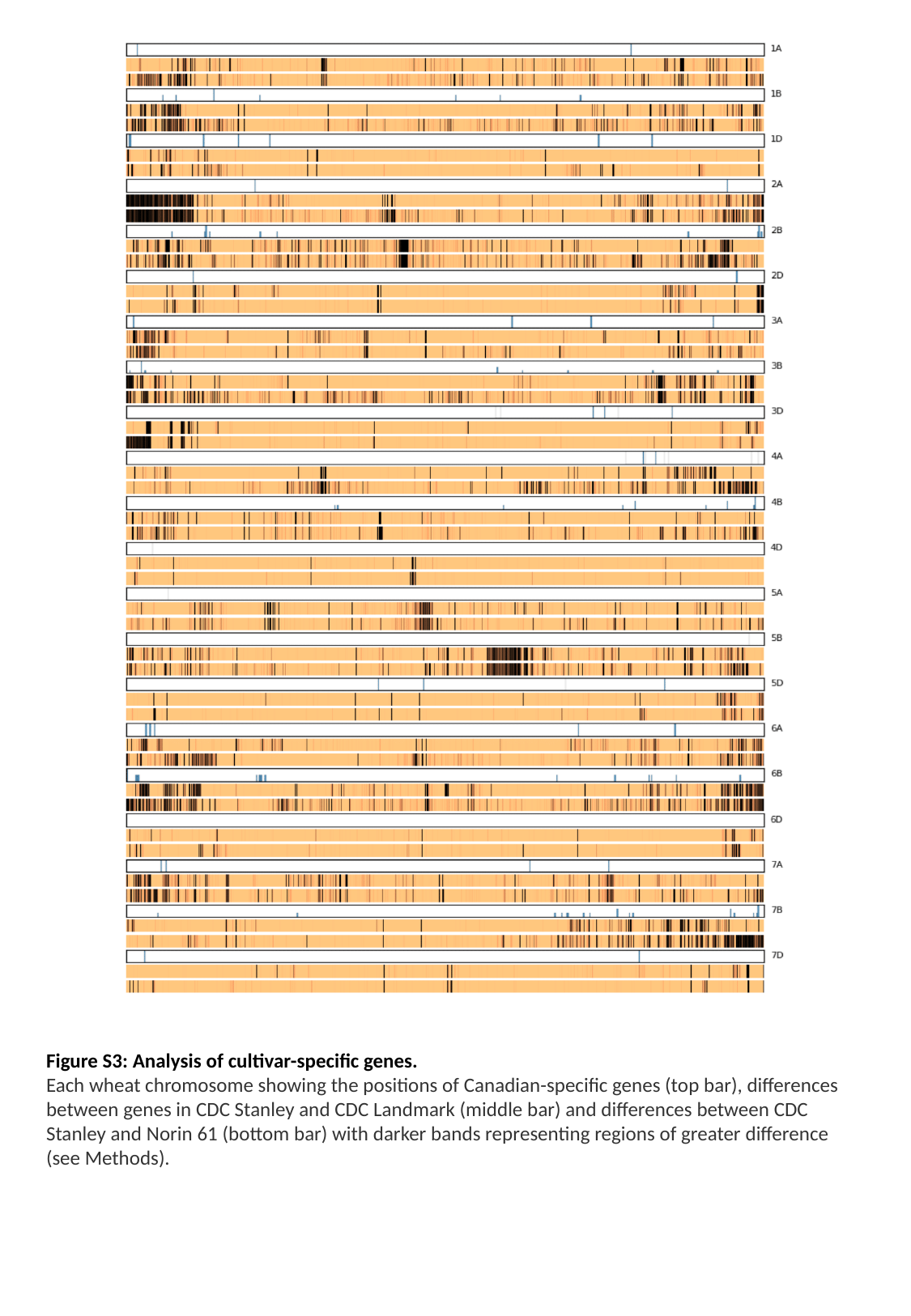

Figure S3: Analysis of cultivar-specific genes.
Each wheat chromosome showing the positions of Canadian-specific genes (top bar), differences between genes in CDC Stanley and CDC Landmark (middle bar) and differences between CDC Stanley and Norin 61 (bottom bar) with darker bands representing regions of greater difference (see Methods).

### Slide 4
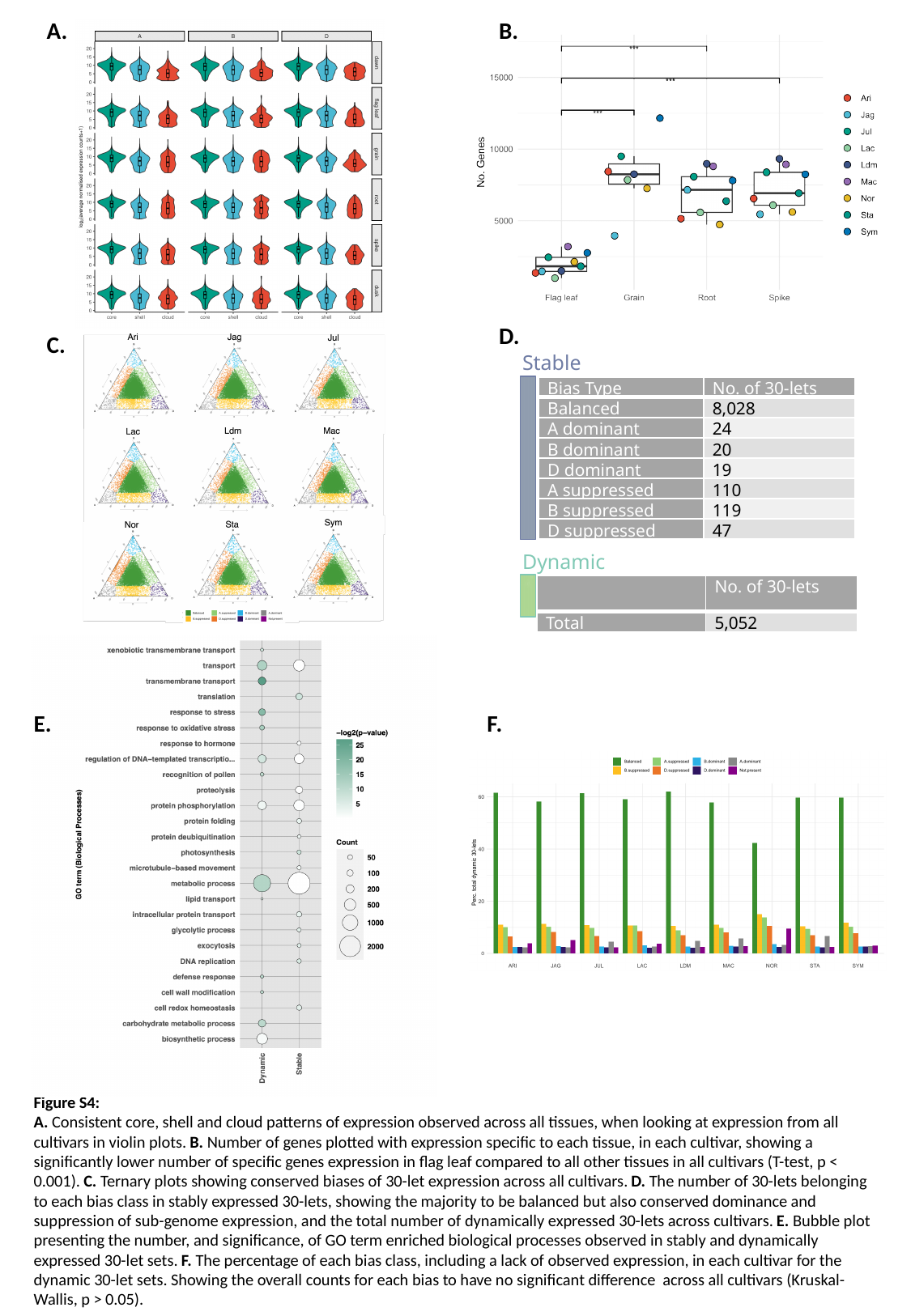

A.
B.
D.
C.
Stable
| Bias Type | No. of 30-lets |
| --- | --- |
| Balanced | 8,028 |
| A dominant | 24 |
| B dominant | 20 |
| D dominant | 19 |
| A suppressed | 110 |
| B suppressed | 119 |
| D suppressed | 47 |
Dynamic
| | No. of 30-lets |
| --- | --- |
| Total | 5,052 |
E.
F.
Figure S4:
A. Consistent core, shell and cloud patterns of expression observed across all tissues, when looking at expression from all cultivars in violin plots. B. Number of genes plotted with expression specific to each tissue, in each cultivar, showing a significantly lower number of specific genes expression in flag leaf compared to all other tissues in all cultivars (T-test, p < 0.001). C. Ternary plots showing conserved biases of 30-let expression across all cultivars. D. The number of 30-lets belonging to each bias class in stably expressed 30-lets, showing the majority to be balanced but also conserved dominance and suppression of sub-genome expression, and the total number of dynamically expressed 30-lets across cultivars. E. Bubble plot presenting the number, and significance, of GO term enriched biological processes observed in stably and dynamically expressed 30-let sets. F. The percentage of each bias class, including a lack of observed expression, in each cultivar for the dynamic 30-let sets. Showing the overall counts for each bias to have no significant difference across all cultivars (Kruskal-Wallis, p > 0.05).

### Slide 5
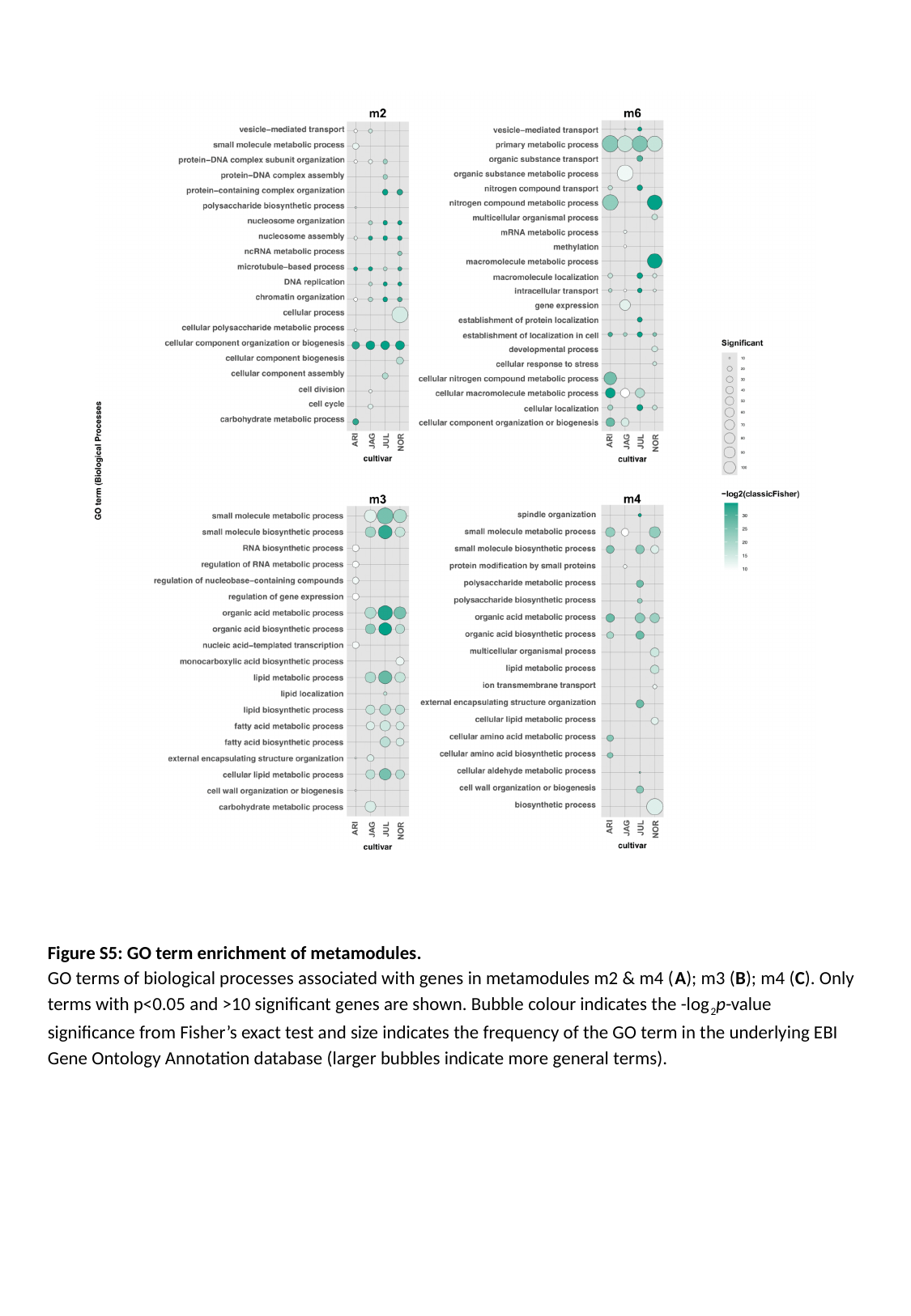

Figure S5: GO term enrichment of metamodules.
GO terms of biological processes associated with genes in metamodules m2 & m4 (A); m3 (B); m4 (C). Only terms with p<0.05 and >10 significant genes are shown. Bubble colour indicates the -log_2_p-value significance from Fisher’s exact test and size indicates the frequency of the GO term in the underlying EBI Gene Ontology Annotation database (larger bubbles indicate more general terms).

### Slide 6
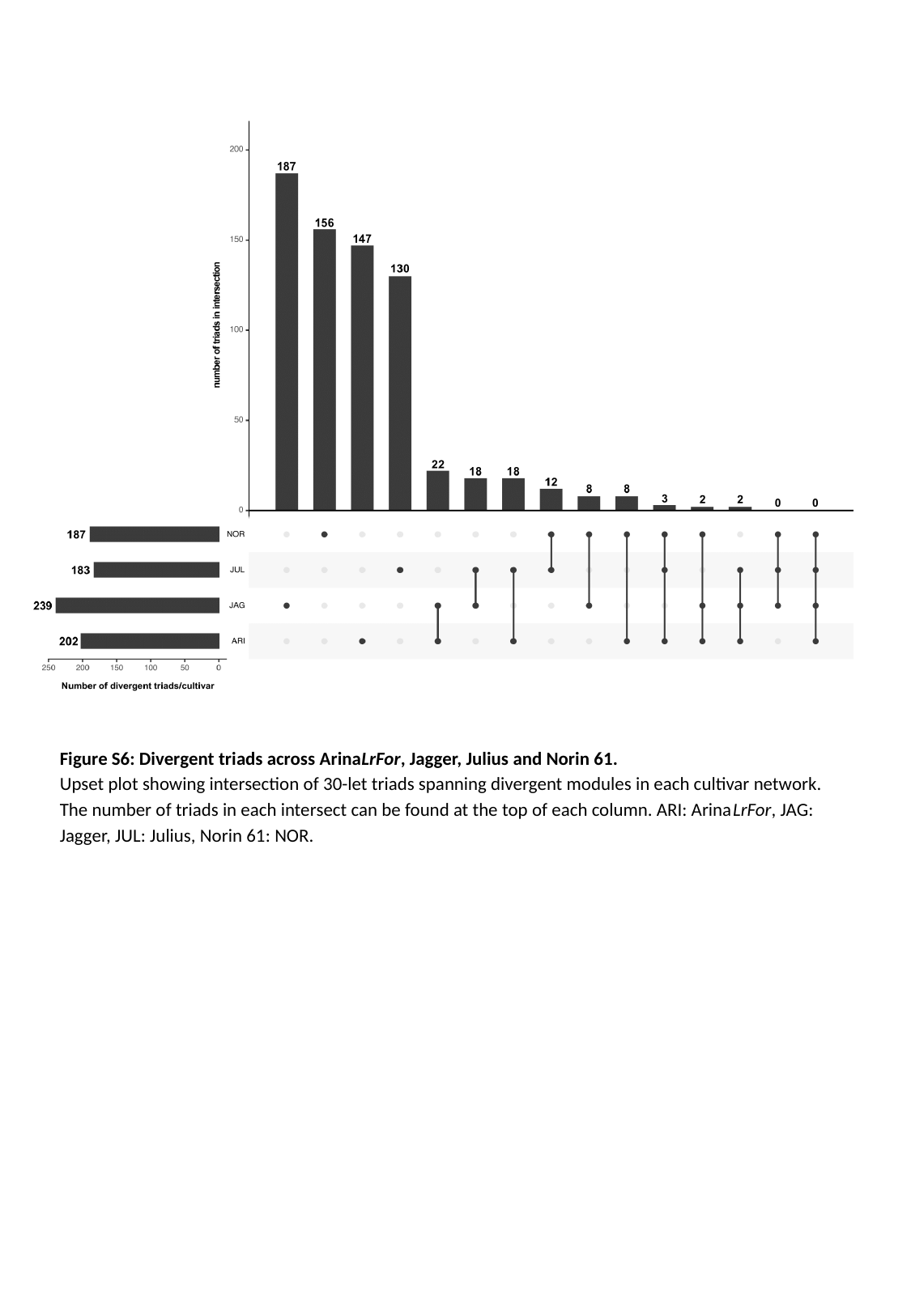

Figure S6: Divergent triads across ArinaLrFor, Jagger, Julius and Norin 61.
Upset plot showing intersection of 30-let triads spanning divergent modules in each cultivar network. The number of triads in each intersect can be found at the top of each column. ARI: ArinaLrFor, JAG: Jagger, JUL: Julius, Norin 61: NOR.

### Slide 7
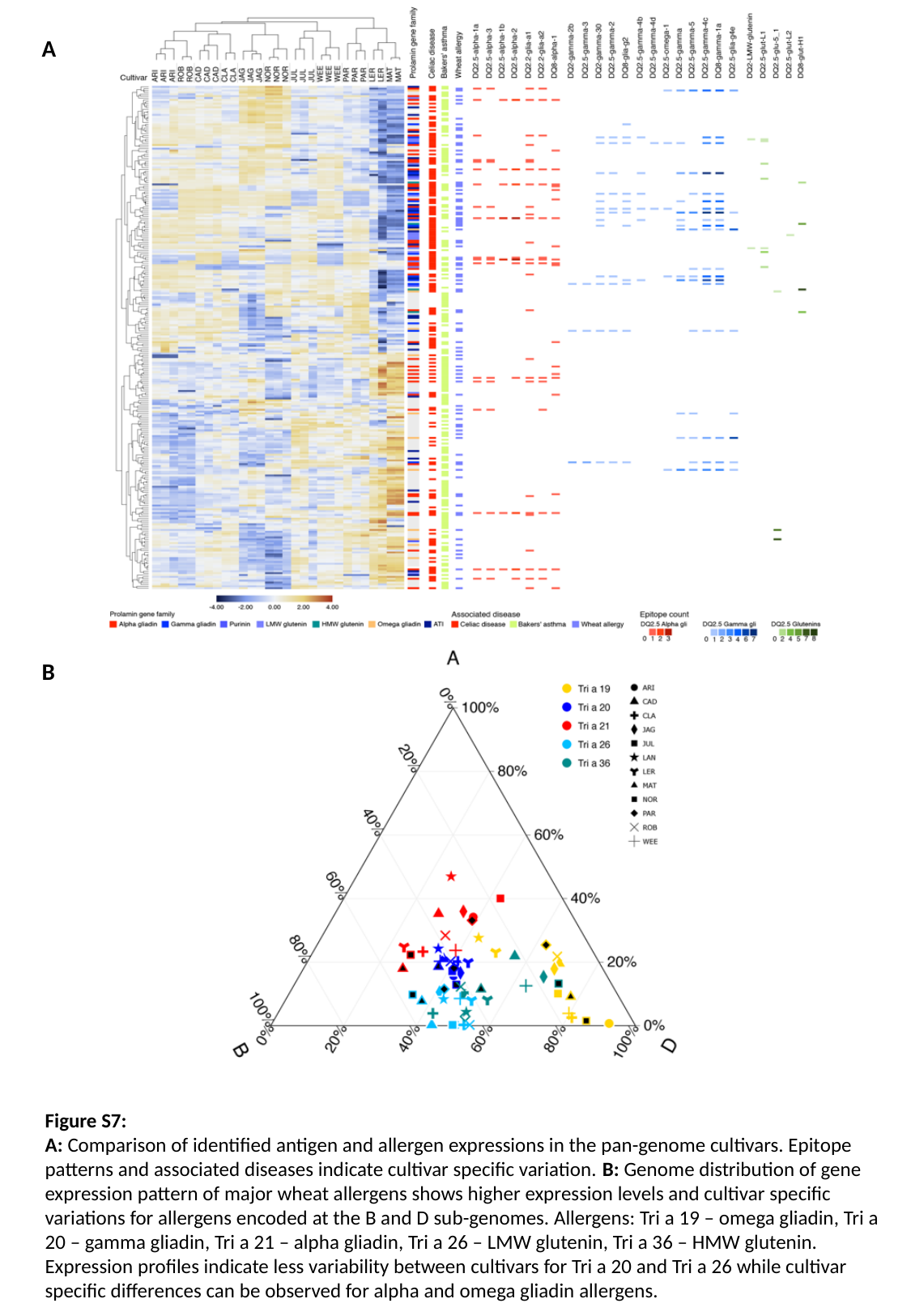

A
B
Figure S7:
A: Comparison of identified antigen and allergen expressions in the pan-genome cultivars. Epitope patterns and associated diseases indicate cultivar specific variation. B: Genome distribution of gene expression pattern of major wheat allergens shows higher expression levels and cultivar specific variations for allergens encoded at the B and D sub-genomes. Allergens: Tri a 19 – omega gliadin, Tri a 20 – gamma gliadin, Tri a 21 – alpha gliadin, Tri a 26 – LMW glutenin, Tri a 36 – HMW glutenin. Expression profiles indicate less variability between cultivars for Tri a 20 and Tri a 26 while cultivar specific differences can be observed for alpha and omega gliadin allergens.

### Slide 8
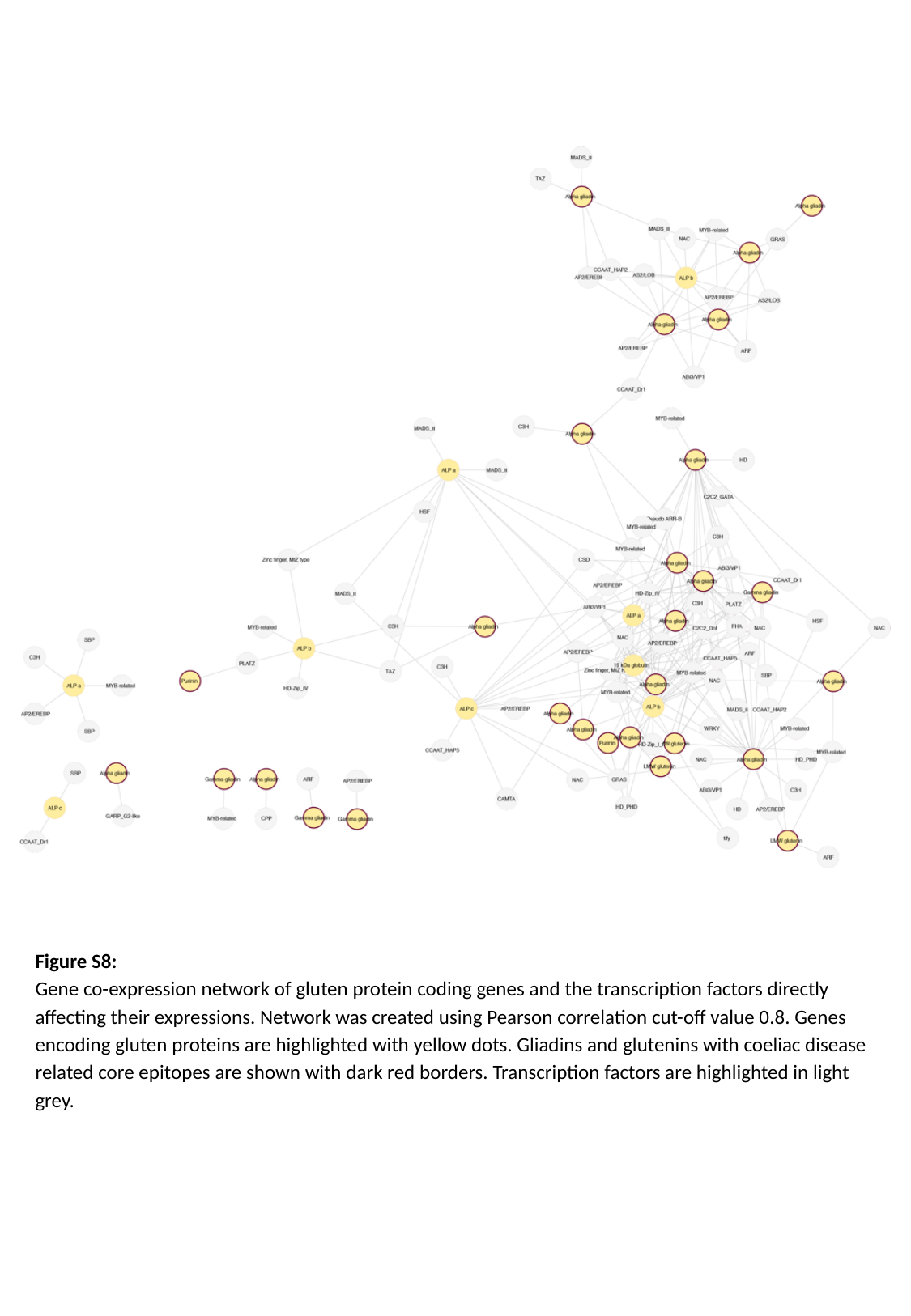

Figure S8:
Gene co-expression network of gluten protein coding genes and the transcription factors directly affecting their expressions. Network was created using Pearson correlation cut-off value 0.8. Genes encoding gluten proteins are highlighted with yellow dots. Gliadins and glutenins with coeliac disease related core epitopes are shown with dark red borders. Transcription factors are highlighted in light grey.

### Slide 9
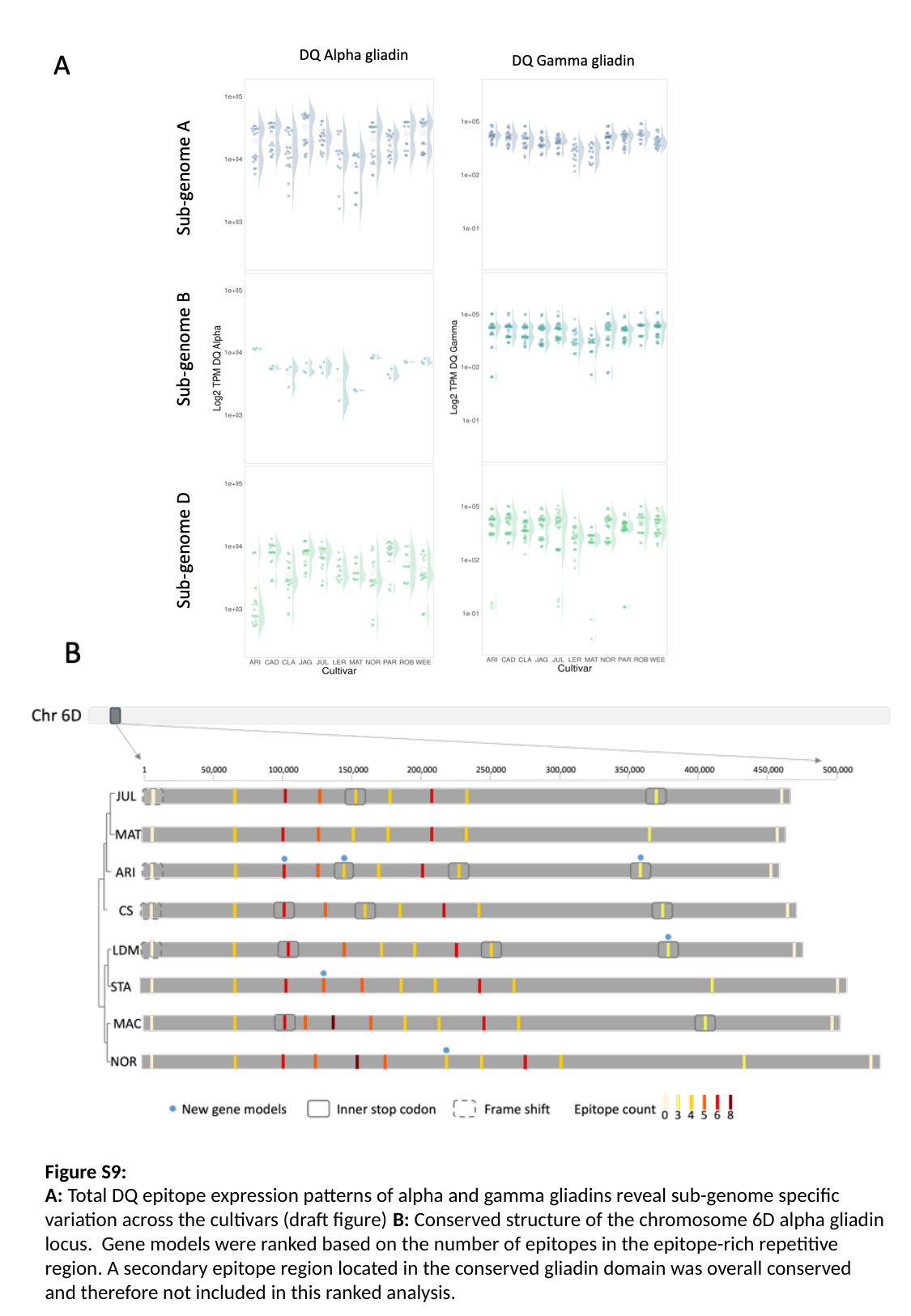

Figure S9:
A: Total DQ epitope expression patterns of alpha and gamma gliadins reveal sub-genome specific variation across the cultivars (draft figure) B: Conserved structure of the chromosome 6D alpha gliadin locus. Gene models were ranked based on the number of epitopes in the epitope-rich repetitive region. A secondary epitope region located in the conserved gliadin domain was overall conserved and therefore not included in this ranked analysis.

### Slide 10
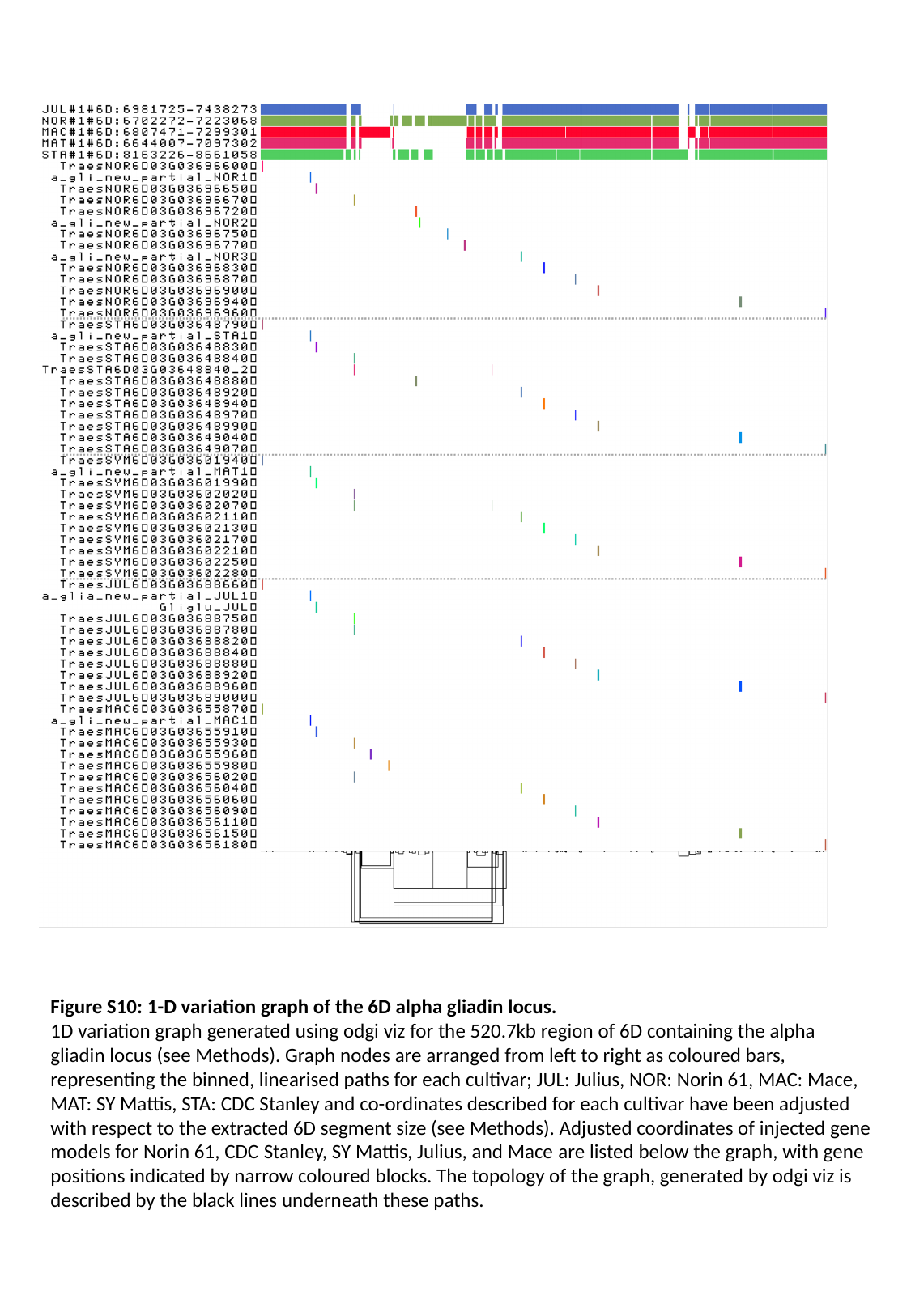

Figure S10: 1-D variation graph of the 6D alpha gliadin locus.
1D variation graph generated using odgi viz for the 520.7kb region of 6D containing the alpha gliadin locus (see Methods). Graph nodes are arranged from left to right as coloured bars, representing the binned, linearised paths for each cultivar; JUL: Julius, NOR: Norin 61, MAC: Mace, MAT: SY Mattis, STA: CDC Stanley and co-ordinates described for each cultivar have been adjusted with respect to the extracted 6D segment size (see Methods). Adjusted coordinates of injected gene models for Norin 61, CDC Stanley, SY Mattis, Julius, and Mace are listed below the graph, with gene positions indicated by narrow coloured blocks. The topology of the graph, generated by odgi viz is described by the black lines underneath these paths.
